# Evaluation of modulators of cAMP-response in terms of their impact on cell cycle and mitochondrial activity of *Leishmania donovani*

**DOI:** 10.1101/2020.01.14.906321

**Authors:** Amrita Saha, Anindita Bhattacharjee, Amit Vij, Pijush K. Das, Arijit Bhattacharya, Arunima Biswas

## Abstract

With the identification of novel cAMP binding effecter molecules in *Trypanosoma*, role of cAMP in kinetopalstida parasites gained an intriguing break through. Despite earlier demonstrations of role of cAMP in survival of *Leishmania* during macrophage infection, there is essential need to specifically clarify involvement of cAMP in various cellular processes in the parasite. In this context, we sought to gain a comprehensive understanding of the effect of cAMPanalogs and cAMP-phosphodiesterase (PDE) inhibitors on proliferation of log phase parasites. Administration of both hydrolysable (8-pCPT-cAMP) and non-hydrolysable analogs (Sp-8-pCPT-cAMPS) of cAMP resulted in significant decrease of *Leishmania* proliferation. Amongst the various PDE inhibitors, etazolate was found to be potently anti-proliferative. BrdU cell proliferation and K/N/F-enumeration microscopic study revealed that both cAMP analogues and selective PDE inhibitors resulted in significant cell cycle arrest at G_1_ phase with reduced S-phase population. Furthermore, careful examination of the flagellar motility patterns revealed significantly reduced coordinated forward flagellar movement of the promastigotes with a concomitant decrease in cellular ATP levels. Alongside, 8-pCPT-cAMP and PDE inhibitors etazolate and trequinsin showed marked reduction in mitochondrial membrane potential. Treatment of etazolate at subcytotoxic concentration to infected macrophages significantly reduced parasite burden and administration of etazolate to *Leishmania*-infected BALB/c mice showed reduced liver and spleen parasite burden. Collectively, these results imply involvement of cAMP in various crucial processes paving the avenue for developing potent anti-leishmanial agent.

**Author Summary:** *Leishmania donovani* is the causative agent of fatal Visceral Leishmaniasis. The current available medications are toxic, expensive and require long term daily administrations. With an aim to develop improved therapeutic, components of cAMP homeostasis, particularly cAMP-phosphodiesteares, has been targeted for *Leishmania* and other kinetoplastid pathogens. cAMP plays diverse roles in functional processes involved in cell division, transition into different stages of the life cycle of *Leishmania* and motility. In this study, the authors found administration of both hydrolysable and non-hydrolysable analogs of cAMP and certain PDE inhibitors resulted in remarkable decrease proliferation with considerable cytopathic impact on promastigotes. The mammalian phosphodiestearse inhibitor etazolate caused significant reduction in parasite load in *L. donovani* infected macrophages and demonstrated considerable reduction of liver and spleen parasite burden in *in vivo* mouse infection model. The study suggested that etazolate, with its slightest impact on mammalian host, can be repurposed for developing effective anti-leishmanials.

## Introduction

Leishmaniasis defines an array of diseases including self-healing cutaneous leishmaniasis and fatal visceral leishmaniasis caused by various species of protozoan parasite *Leishmania*. Till date, the disease is considered as a neglected tropical disease and urgently requires new therapeutic interventions. For leishmaniasis, pentavalent antimonials have remained the standard medication till recent decades, but drug resistance is becoming an increasing problem in the disease prone areas [1]. Although a number of new compounds, such as miltefosine or amphothericin B, have been developed over the last few years [2–6], effective, safe and cost-efficient chemotherapy of leishmaniases still remains elusive. It has been currently postulated that cAMP plays an important role in various cellular processes linked with survival and virulence of kinetopalstida parasites [7,8]. In *Trypanosoma brucei* detailed studies revealed that hydrolysed product of cAMP as well as various isoenzymes of cyclic nucleotide phosphodiesterases (PDE) and adenylatecyclase (AC) have an important role in parasite survival, virulence and cell-cell communications [7,8]. Moreover, the isoform PDEC from *T. cruzi* was shown to be involved in osmoregulation and the potential of it to be a drug target was validated by high throughput screening of a series of PDE inhibitors [9], which led to identification of tetrahydrophthalazinone compound A (Cpd A) [7]. These observations underpin the possibilities of cAMP pathway as a candidate target for drug development [9].

*Leishmania* differs in terms of its repertoire of cAMP-pathway components from *T. brucei* as the number of AC genes is considerably less. Additionally it encodes one cytosolic heme-containing AC which is absent in *Trypansoma*, indicating presence of subcellular microdomain specific cAMP signalling in this parasite [10,11]. Elevation of cAMP was found to be a prerequisite for differentiation condition induced cell cycle arrest (G1) and adaptation against oxidative stress encountered during early macrophage invasion [12,13]. Moreover, the crucial role played by both exogenous and endogenous cAMP in chemotaxis by sensing chemical cues and inducing flagellar wave reversal [14] suggest the existence of definitive cAMP regulated mechanisms in the parasite. Similar to other kinetoplastida parasites, *Leishmania* encodes at least 7 different cyclic nucleotide targeted phosphodiesterases (PDE). X-ray crystallography structure of *L. major* PDEB1 and B2 has been determined and it’s superposition with human PDEs revealed presence of a unique sub pocket nearby inhibitor binding sites [15], widening the scope of selective drug designing. The differentially regulated *L. donovani* PDEA was characterized to be a high K_M_ cytosolic PDE that regulates cytosolic cAMP pool and modulate expression of anti-oxidant genes [16]. The other cytosolic PDE, PDED was found to be a moderate K_M_ enzyme that interact with PKA-catalytic subunit like proteins to regulate cAMP level [17]. Early studies showed that three human PDE inhibitors (etazolate, dipyridamole and trequinsin) inhibit the proliferation of *L. major* promastigotes and *L. infantum* amastigotes with IC_50_ values in the range of 30–100 μM [18]. Moreover, dipyridamole, etazolate and trequinsin were observed to be potent inhibitors of *L. donovani* PDEA [16] whereas rolipram was observed to inhibit PDED amongst all the mammalian PDE inhibitors [17]. A robust PDE inhibitor library, has been reported emphasizing PDE as exploitable targets for intervention [19]. However, a comprehensive profile for the impact of PDE inhibitors on various cellular processes in *Leishmania* is still lacking.

Although several cAMP and cAMP pathway associated phenotypes has been identified in *Leishmania* any functional effector for cAMP is yet to be identified in this parasite. With an intention to assess the potential of cAMP pathway as target for chemotherapeutic intervention, in the present study, we intended to screen various cAMP analogues and PDE inhibitors. It was observed that hydrolysis-resistant cell-permeable cAMP analogues possess potent anti-proliferative effect and show G_0_/G_1_ phase arrest indicating that unlike in *Trypanosoma*, where products of cAMP hydrolysis seemed vital for parasite transformation, in *Leishmania* cAMP itself could induce these effects. Moreover, mammalian PDE4 inhibitor rolipram and etazolate and PDE5 inhibitor dipyridamole showed more anti-proliferative effect and cell cycle arrest compared to the other well-known mammalian PDE inhibitors. Such effects were also found to be associated with slow and sluggish movement of *Leishmania* promastigotes, a change in ATP level and a striking change in mitochondrial membrane potential, reinforcing the need for further exploration of cAMP pathway in the parasite for drug development. Efficacy of etazolate in mouse model of VL and apparent unresponsiveness of mammalian cells against this PDE inhibitor validate etazolate as a prospective repurposed drug against *Leishmania*.

## 2. Results

### 2.1. Effects of cAMP analogues and PDE inhibitors on *L. donovani*promastigotes

The effect of various cAMP analogues and PDE inhibitors on the growth of log-phase promastigotes of *L. donovani* was measured by proliferation assays where parasites were grown in presence of various concentrations of these compounds. Intracellular cAMP concentration was modulated by treatment with one hydrolysable analogue, 8-(4-chlorophenylthio)adenosine 3’,5’-cyclic monophosphate (8-pCPTcAMP), and one non-hydrolysable analogue, 8-(4-chlorophenylthio)adenosine-3’,5’-cyclic monophosphorothioate, Sp-isomer (Sp-8-pCPTcAMPS). Both the analogues significantly affected growth with EC_50(growth)_ of 218.22 μM and 189.78 μM respectively (Table I). On the flip-side, non-hydrolysable cGMP-analogue, 8-(4-chlorophenylthio)-guanosine 3’,5’-cyclic monophosphate (8-pCPT-cGMP) failed to show any anti-proliferative effect on the parasites. Inhibitors previously determined to inhibit Leishmania-PDEs including EHNA, etazolate, dipyridamole, zaprinst, rolipram and trequinsin, were profiled for prospective antiprolifearative activity. Dose-response curves were obtained by calculating the percentage proliferation against untreated population after 4 days of growth to determine EC_50(growth)_. Etazolate was found to be potently anti-proliferative with EC_50(growth)_ of 24.1 μM (Table 1). Dipyridamole and rolipram showed anti-proliferative effects to lesser extents (EC_50(growth)_ of 61.3 μM and 29.04 μM respectively). Trequinsin on the other hand showed comparatively lower anti-proliferative effect with EC_50(growth)_ of 79.6 μM. Rest of the PDE inhibitors tested showed little effect on cell proliferation (Table 1).

**Table-1.**
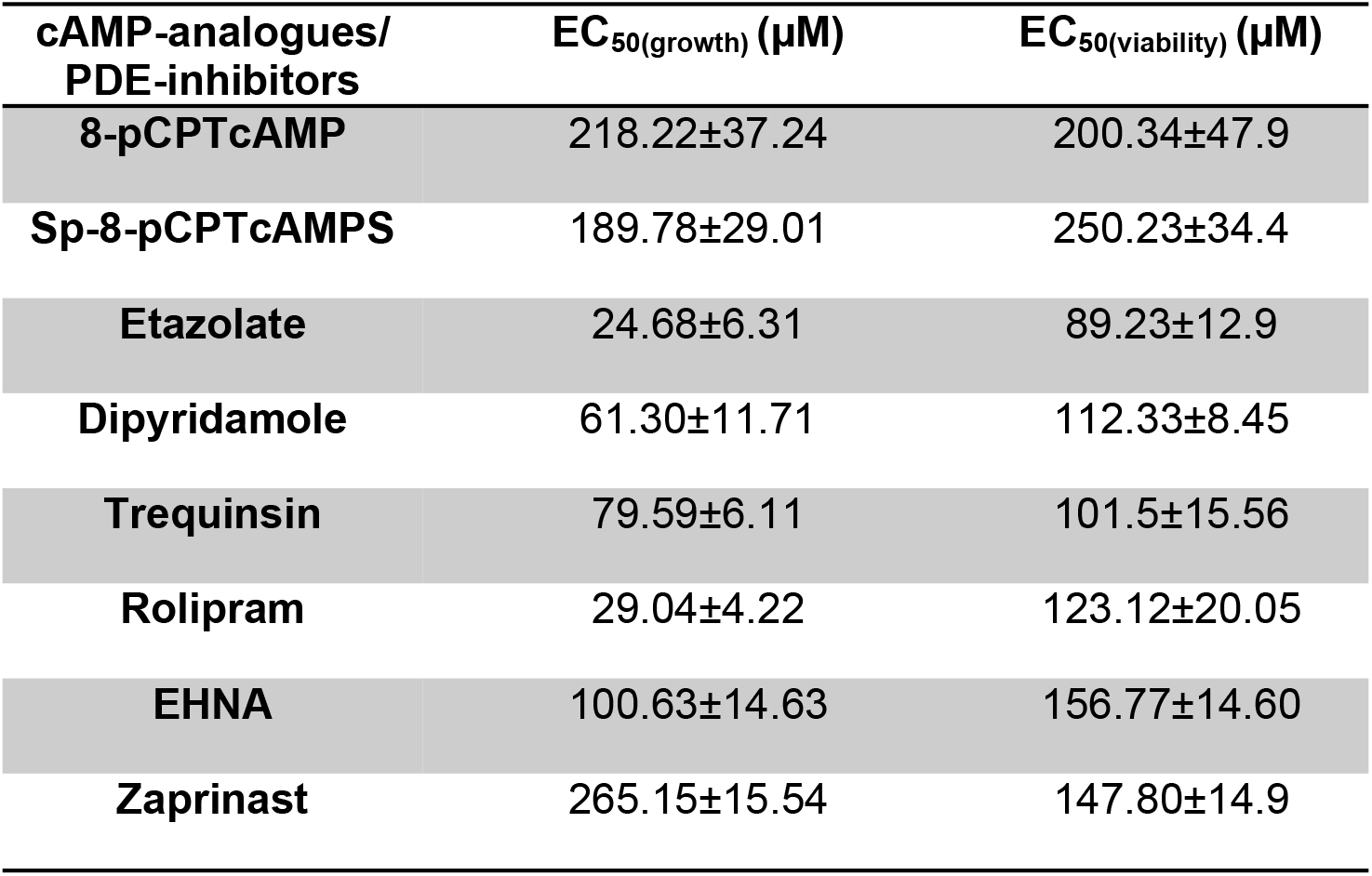
Effect of cAMP analogues and PDE=inhibitors on *L. donovani*promastigotes. EC_50(growth)_ was calculated after allowing the promastigotes to grow in presence of analogues/ inhibitors for 96h. EC_50(viability)_ was tested by MTT based vital assay after exposing the promastigotes to the analogues and inhibitors for 24h. Data are representative of at least three independent experiments.

Since cAMP analogues and PDE inhibitors showed anti-proliferative effects on the parasite, we were interested to introspect whether those impart any effect on parasite viability by determining EC_50(viability)_ by MTT assay after 24 h of exposure. 8-pCPTcAMP and 8-Sp-pCPTcAMPS significantly decreased the parasite viability with EC_50(viability)_ 200.34 μM and 250.23 μM respectively (Figure S1A and S1B; Table 1). 8-pCPTcGMP apparently did not affect parasite viability (data not shown). EC_50(viability)_ values were also determined from the dose responsive curves of various PDE inhibitors (Figure S1). Etazolate, with an EC_50(viability)_ of 89.23 μM, appeared as the most promising anti-proliferative of the inhibitors tested. Dipyridamole, trequinsin and rolipram showed EC_50_ values of 112.33, 101.5 and 123.12 μM respectively (Table 1, Figure S1C-S1F). All the other PDE inhibitors showed higher EC_50(viability)_ values (Table 1). Collectively, these results suggest that cAMP analogues and PDE inhibitors affects parasite viability at higher doses.

### 2.2. Effect of cAMP analogues and PDE inhibitors on the cell cycle

To assess the effect of cAMP analogues and PDE inhibitors on the cell cycle, promastigotes were treated with increasing concentrations (10, 50 and 100 μM) for 12 h followed by 2 h incubation with BrdU and the percentage BrdU incorporation was studied by BrdU cell proliferation assay. 8-pCPTcAMP and 8-Sp-pCPTcAMPS treatment at 100 μM concentration significantly decreased the number of BrdU-positive cells (34.2% and 31.64% respectively compared to 82.3% in control) (Figure 1A), indicating fewer cells progressing into the S-phase of the cell cycle. Treatment with PDE inhibitors etazolate, dipyridamole, trequinsin and rolipram also decreased the BrdU uptake significantly at 50 μM (24.13%, 28.66%, 31.86% and 26.56% compared to untreated control of 82.38%) whereas, PDE inhibitors EHNA and zaprinast showed only 73.76% and 68.5% positives at 50 μM respectively (Figure 1B). To further introspect the impact on cellular proliferation, progression of cell division was determined by scoring of the number, dimension and position of the nucleus (N), kinetoplast (K) and flagellum (F) in cAMP analogues and PDE inhibitor-treated cells after labelling with DAPI. Cells with one of each of these organelles - 1 kinetoplast/1 nucleus/1 flagella (1N/1K/1F) representing the G0/G1 population enter S phase and duplicate their DNA content (1N*/1K*/1F cells, where * denotes duplicated DNA). Initiation of the growth of the second flagellum then begins (1N^*^/1K^*^/2F cells), where upon segregation of the two daughter kinetoplasts is initiated and 2K1N2F represents S phase followed by nuclear segregation (2N/2K/2F cells) which represents G2/M phase population [20]. Treatment with 8-pCPTcAMP and 8-Sp-pCPTcAMPS resulted in a marked increase in the 1K/1N/1F population, from 54.8 to 75.93% and 78.03% respectively after 12 h, which were associated with concomitant decrease in the 2K/2N/2F and 2K/1N/2F populations (Figures 1C and 1E). Exposure to PDE inhibitors etazolate, rolipram, dipyridamole and trequinsin resulted in a significant increase in the 1N/1K/1F cells (73.43% ± 4.3%, 75.16% ± 5.1%, 75% ± 4.9% and 71.5% ± 6.4% compared to 51.53% ± 4.7% for control after 12 h) with equivalent decrease in 2N/2K/1F cells and 2N/2K/2F cells (Figures 1D and 1E). Treatment with PDE inhibitors zaprinast and EHNA (Figure 1D) showed K/N/F profiles comparable to that of the untreated cells. Taken together, these results suggest a probable G1 arrest of *Leishmania* promastigotes when they were treated with cAMP analogues and PDE inhibitors.

**Fig.1.**
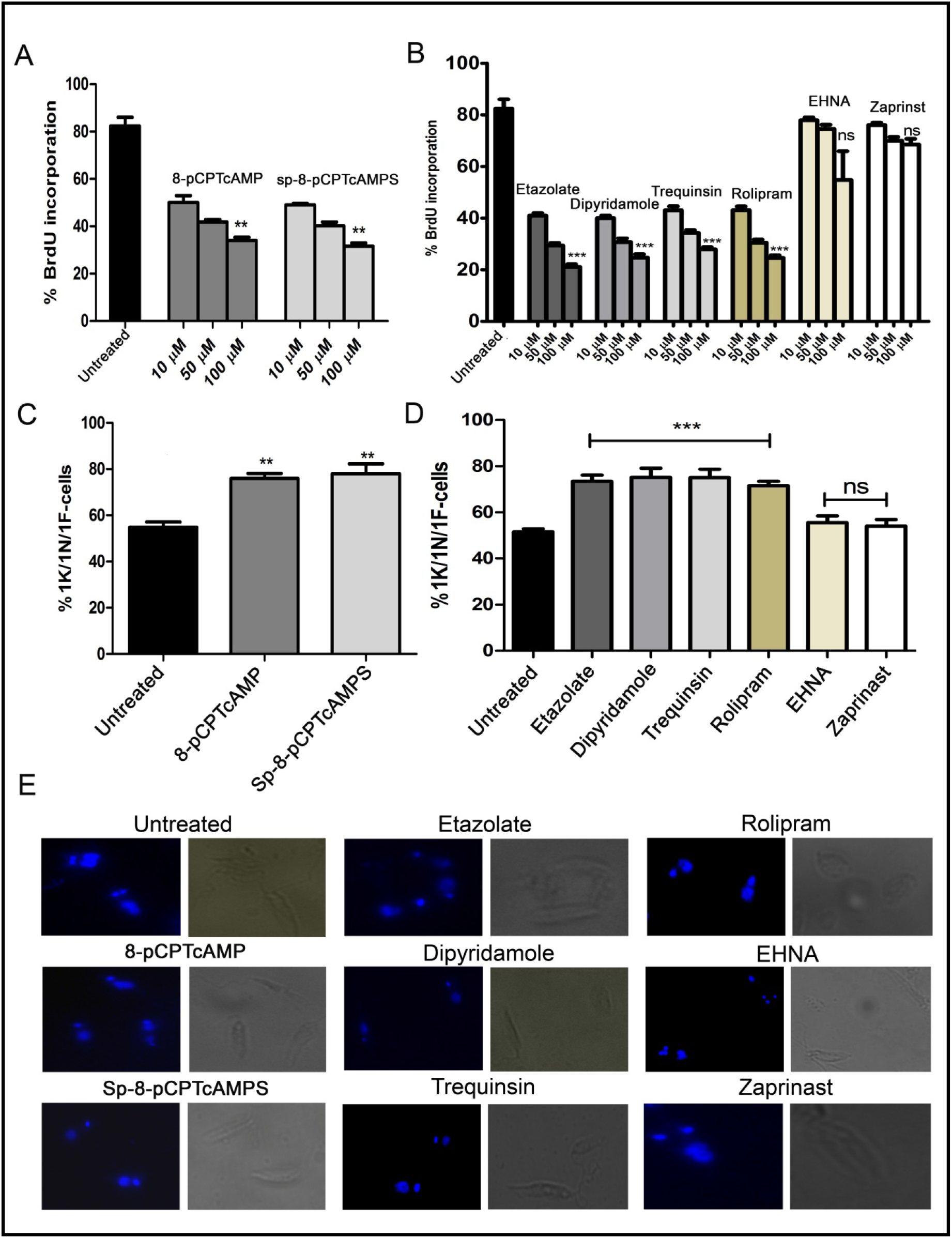
Effect of cAMP analogues and PDE inhibitors on the cell cycle. (A and B) Log phase promastigotes were pretreated with various concentrations of cAMP analogues (A) and PDE inhibitors (B) for 6 h followed by BrdU incorporation assay. (C and D) Log phase promastigotes were treated with cAMP analogues (100 μM) (C) and PDE inhibitors (50 μM) (D) for 12 h followed by DAPI staining and 1K/1N/1F cells were scored for each of the treatment from at least 50 fields. Representative confocal microscopic images of the above experiment and Phase-contrast images of each treatment are shown separately (E). Data are representative of three independent experiments. ***P<0.001; **P<0.01; ns not significant, two tailed unpaired t-test.

To confirm the G1 phase cell cycle arrest, G_1_-phase-synchronized promastigotes were introduced into fresh medium and the cell cycle profiles of cAMP analogue-treated and PDE inhibitor-treated populations were studied for cellular DNA estimation by propidium iodide based FACS analysis. After exposing the G1 synchronized cells to 8-pCPTcAMP and Sp-8-pCPT-cAMPS (10 μM, 50 μM 100 μM), parasites were tracked over a period of 8, 16 and 24 h for cell cycle progression in G_1_, S and G_2_/M phase (Figure 2A and Table S2). Treatment of the cells with 8-pCPT-cAMP reduced the S phase fraction indicating a potent role of 8-pCPTcAMP induced cell cycle arrest at G_1_ phase, which also indicated by an increase in G_1_ population (Figure 2B and Table S2). Similarly, Sp-8-pCPT-cAMPS reduced the percentages of S phase cells significantly and increased G1 phase population (Figure 2C and Table S2). For treatment with PDE inhibitors etazolate, dipyridamole and trequinsin, inhibitory toward *Leishmania* PDEA and PDEB [18], and rolipram, inhibitory toward *Leishmania* PDED [13], showed significant G_1_ arrest after 24 h while for inhibitors like EHNA and cGMP-PDE specific inhibitor zaprinst, no apparent G1 arrest was observed. Etazolate, dipyridamole, rolipram and trequinsin resulted significantly reduced S phase cell population at 100 μM respectively compared to untreated population (Table-S2 and Figures 2D to I). These results indicate that chemical modulation of intracellular cAMP can affect cell cycle progression.

**Fig.2.**
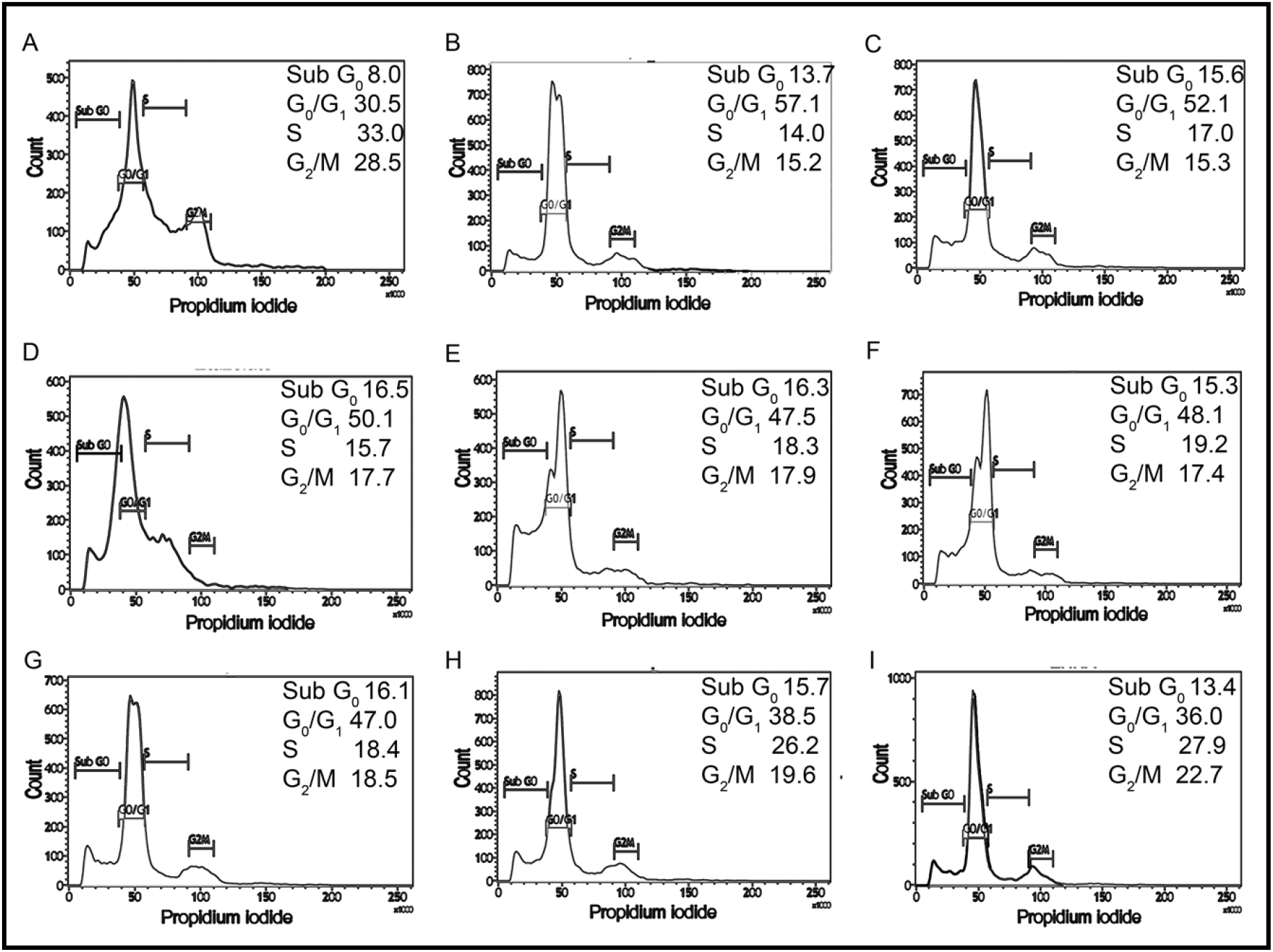
Impact of cAMP analogues and PDE inhibitors on cell cycle progression of *L. donovani* promastigotes.G1-synchronized log phase promastigote populations, either untreated (A) or treated with 8-pCPTcAMP (100 μM) (B), Sp-8-pCPTcAMPS (100 μM) (C); etazolate (D), dipyridamole (E), trequinsin (F), rolipram (G), EHNA (H) or zaprinast(I) (50 μM each) for 8h, 16h and 24h, were analysed for cell cycle progression by propidium iodide staining based FACS analysis (Table S1). Representative histograms from 24h data are presented.

### 2.3. Modulation of parasite motility and ATP level by dose-dependent treatment of cAMP analogues and PDE inhibitors is linked to mitochondrial activity

Since cAMP has a role in flagellar motility of *Leishmania* and social motility of *Trypanosoma* [21], an examination of their flagellar motility patterns revealed that at a concentration of 100 μM both 8-pCPTcAMP and Sp-8-pCPTcAMPS affected the coordinated progressive flagellar movement or forward swimming motility of the parasites significantly (Figure 3A). Parasites were also significantly impaired for progressive flagellar movement in presence of etazolate, dipyridamole and rolipram (Figure 3A). On the other hand, EHNA and zaprinast, did not affect forward flagellar movement (Figure 3A). Flagellar length and ratio of cell body-flagellar length have been associated with flagellar motility [22]. Average flagellar and cell body length were measured from 50 cells each from three independent populations of cells treated with the inhibitors. Apart from minor increase for dipyridamole treatment, none of the cAMP analogues and PDE inhibitors inducted alteration in flagellar length (Figure 3B). Since, one of the factors effecting flagellar motility is intracellular ATP generation, we determined the cellular ATP levels by ATP bio-luminescent assay after treating the parasites with cAMP analogues and PDE inhibitors. Treatment of parasites with various concentrations of cAMP analogues and PDE inhibitors showed that at higher doses of cell permeable cAMP analogues; and PDE inhibitors etazolate, dipyridamole, trequinsin and rolipram significantly reduced the ATP levels (64.8% ± 5.8%, 61.0% ± 5.5%, 49.7% ± 4.1% and 54.7% ± 5.1% respectively) at a concentration of 50 μM whereas 8-pCPTcAMP and Sp-8-pCPTcAMPS caused lesser effect (31.9% ± 2.8% and 26.0% ± 2.2% of reduction respectively) at 100 μM concentration (Figures 3C and 3D). Taking together these results suggest higher concentrations of 8-pCPTcAMP, Sp-8-pCPTcAMPS, rolipram, dipyridamole, etazolate significantly restricts flagellar movement and parasite motility with a concomitant decrease of cellular ATP levels.

**Fig.3.**
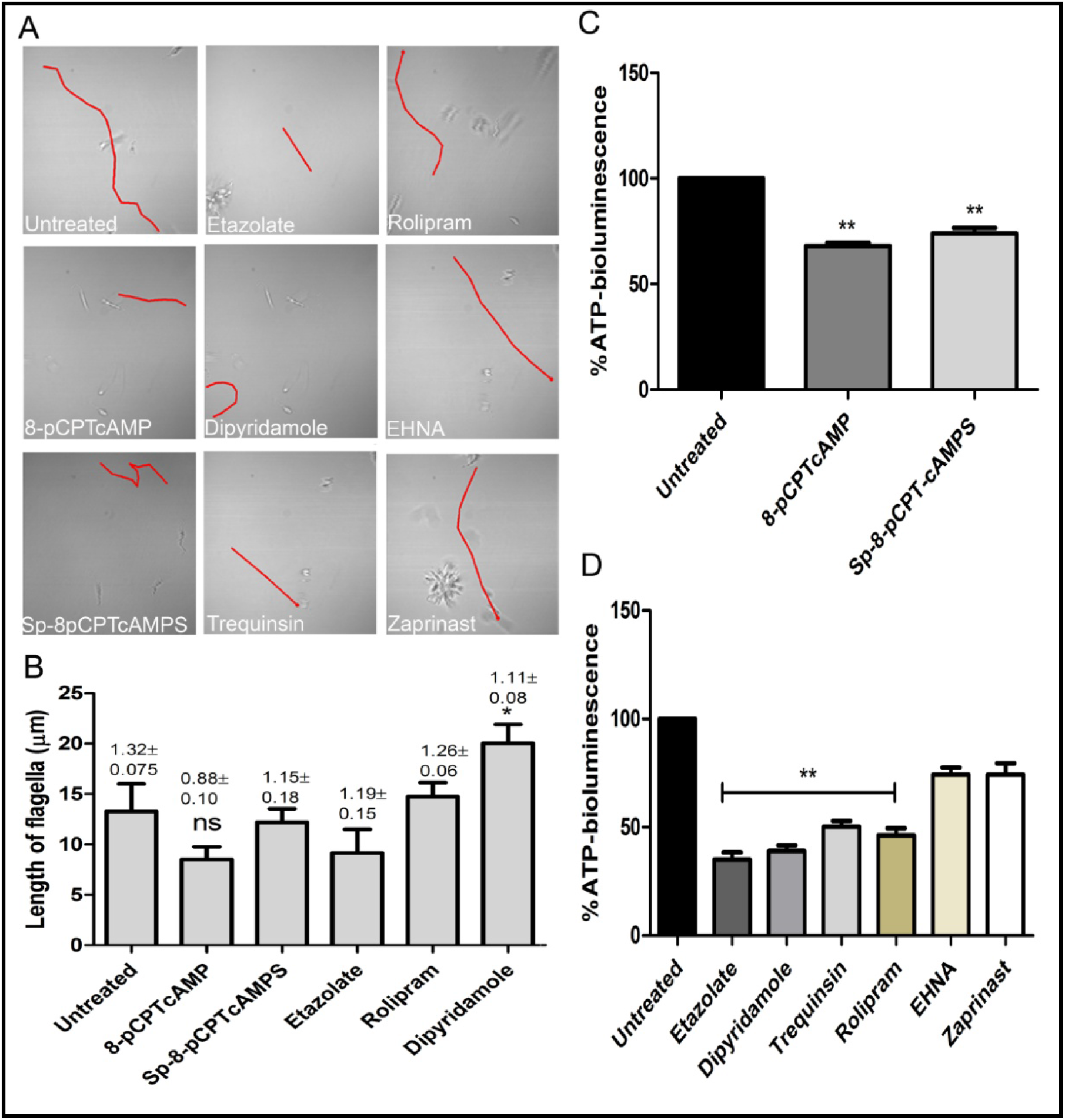
Inhibition of parasite motility and depletion of ATP-level upon administration of specific cAMP-analogues and PDE inhibitors. *Leishmania*promastigotes were treated with various cAMP-analogues (100 μM) and PDE inhibitors (50 μM) and the flagellar motility patterns of both the untreated and treated parasites were studied under the bright field of confocal microscope (A). From the same set of experiments, flagellar length and ratio of flagellar length and cell body length was calculated (shown above each bar) (B). Levels of cellular ATP generation were studied after treating the parasites with cAMP-analogues (C) and PDE inhibitors (D) by ATP bio-luminescence assay. Results are representative of three individual experiments, and the error bars represent mean ± SEM. **P<0.01; *P<0.05; two tailed paired t-test.

Since the ATP levels of cAMP analogue-treated and PDE inhibitor-treated parasites were depleted, the impact of these on mitochondrial function was further elucidated by scoring mitochondrial membrane using JC-1. 8-pCPTcAMP induced maximum reduction in mitochondrial membrane potential compared to control cells reduction (26.1% ± 1.9%) whereas Sp-8-pCPTcAMPS induced moderate effect (10.2% ± 0.9% reduction) at 100 μM (Figures 4A - 4C). Amongst the PDE inhibitors, etazolate and dipyridamole showed maximum reduction (26.4% ± 2.2% and 22.5% ± 1.9% respectively) whereas trequinsin and rolipram showed moderate effect. EHNA and zaprinast did not show significant effect.

**Fig.4.**
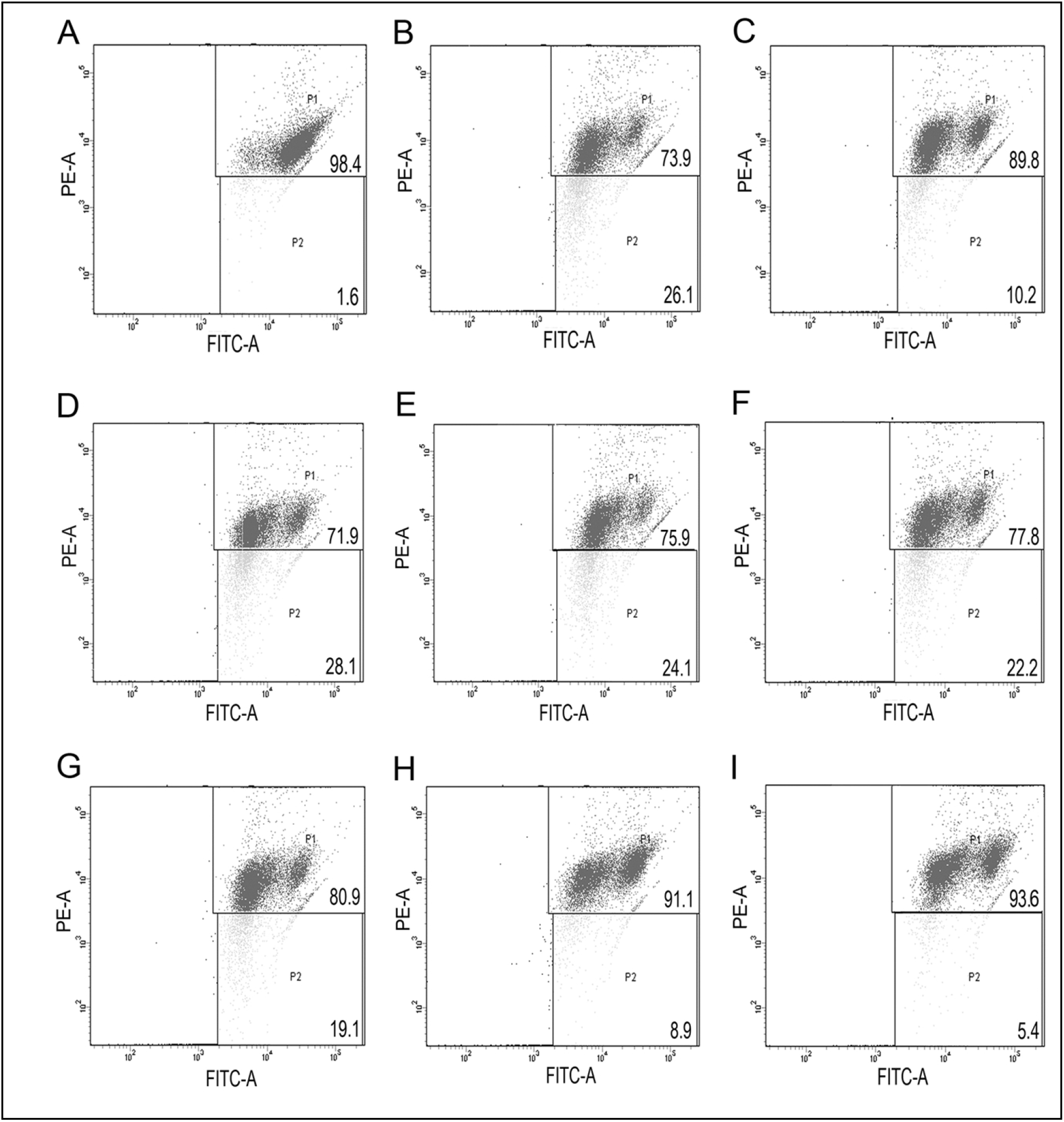
Modulation of mitochondrial membrane potential by cAMP analogues and PDE-inhibitors. Log phase promastigotes were incubated without (A) or with 8-pCPTcAMP (100 μM) (B), Sp-8-pCPTcAMPS (100 μM) (C); etazolate (D), dipyridamole (E), trequinsin (F), rolipram (G), EHNA (H) or zaprinast (I) (50 μM each) for 12h and mitochondrial membrane potential in the parasites were analysed by FACS after treating them with using JC-1, a mitochondrial vital dye. Representative scattered plots for three independent experiments are shown.

### 2.4. Etazolate mitigates intra-macrophage survival of parasites

Since etazolate showed maximum effect on cell cycle progression and caused mitochondrial membrane depolarization, the inhibitor was further explored as anti-leishmanial by analysing its impact on parasite multiplication within RAW 264.7 macrophages. Etazolate apparently had no effect on viability of macrophage, as determined by MTT assay and gross cell morphology analysis as visualized by microscopy up to 200 μM (data not shown). Etazolate treatment resulted in significant reduction in the number of parasites within macrophages with a maximum effect (75.0 % ± 6.7% reduction) observed at 100 μM concentration (Figures 5A and 5B).

**Fig.5.**
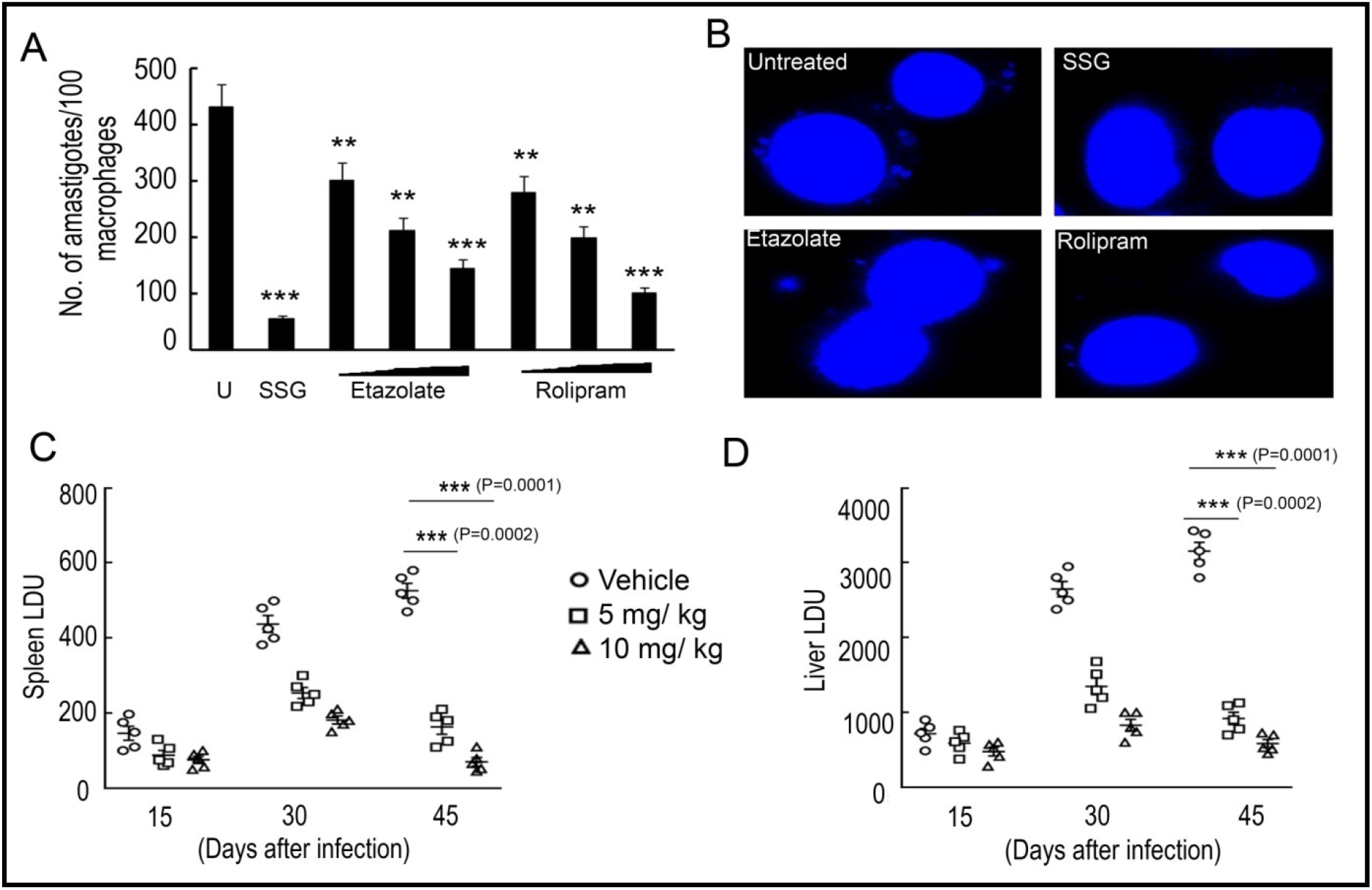
Effect of etazolate on parasite survival. Macrophages were pre-incubated with either increasing concentrations of rolipram and etazolate (1, 10 and 20 μM) or sodium stibogluconate (100 μg/ ml) or left untreated for 30 min, followed by infection with *L. donovani* for another 72 h. The number of parasites per macrophage was scored for untreated (U) and treated systems by DAPI staining (A). Representative microscopic image of infected macrophages either untreated or treated with sodium stibogluconate (100 μg/ ml), rolipram (20 μM) or etazolate (20 μM) are shown (B). BALB/c mice were infected with 10^7^ *L. donovani*promastigotes and treated etazolate (0-10mg/kg/day). Etazolate was administered i.p. twice a week for 6 weeks starting at 1 week post-infection. PBS was used as vehicle control. The parasite burden in liver and spleen was determined 6 weeks after infection and expressed as Leishman–Donovan Units (LDU). Disease progression was determined in spleen (C) and liver (D) of mice that received etazolate at either 5 or 10 mg/kg/day twice weekly and parasite burden expressed as LDU at 2, 4 and 6 week post-infection (C and D). Results are representative of three individual experiments, and the data represent mean ± SD. ***P<0.001, **P<0.01, *P<0.05; unpaired two tailed t test.

Etazolate was tested in BALB/c mice to test its efficacy to clear parasitic burden from spleen and liver in infected groups. A dose of 5 mg/kg/day and 10 mg/kg/day was administered orally over a period of 10 days following 1 month of *Leishmania* infection in mice. Spleen and liver were removed from treated and untreated 45 day infected mice, and multiple impression smears were prepared and stained with Giemsa. Spleen or liver parasite burdens, expressed as Leishman-Donovan units (LDU), were calculated as the number of parasites per 1000 nucleated cells/ organ weight (in grams). A dose-related inhibition was noted at a dose of 5 mg/kg/day and 10 mg/kg/day with great reduction of liver and spleen parasite burden by 70.7 % ± 6.9% and 79.5 % ± 7.2% compared to untreated infected mice (Figures 5C and 5D). These results highlight etazolate as candidate anti-leishmanial.

## Discussion

Leishmaniasis represents a significant global burden and poses a great challenge to drug discovery and delivery, because of the intracellular nature and disseminated locations of the parasite. Most of the current modes of treatment available till date are not only toxic but also do not completely cure, i.e., eliminate the parasite, from infected individuals [20]. The problem of increased chemoresistance towards the current set of available drugs is further complicating the treatment for this disease [22,23]. Because treatment is a growing problem, the development of new medicines that can replace or complement the current set of available therapeutic compounds is necessary.

cAMP has been emerging as one of the major modulator of cytological events in kinetoplastida parasites [8]. Despite the identification of several cyclic nucleotide binding effector molecules there is severe deficit of comprehensive knowledge on cyclic nucleotide signalling from receptor to effector till date. The candidate regulatory subunits and cyclic nucleotide binding domains binds cAMP/ cGMP with extremely high and beyond physiologically permissible Kd indicating almost absence of effective interaction and to perform as a typical effector under physiological condition [15] is not feasible. Bachmaier et al. [24], recently reported binding of nucleotides such as 7-deazapurine derivatives to PKA-regulatory subunit of *T. brucei* instead of canonical cAMP or cGMP. While studying resistance against PDE inhibitor cpdA and cpdB, a novel class of cAMP effectors, CARP was identified in *T. brucei* [25], indicating existence of non-canonical cAMP effectors in kinetoplastida parasites. CARP and its orthologues [25] are conserved among several kineoplastida parasites. On the receptor/cAMP generator side of the pathway, an oxygen stimulated soluble adenylate cyclase has been identified, however since this gene is absent in *Trypanosoma*, this enzyme probably contributes to some cAMP-dependent function restricted to *Leishmania*. The activation of conserved receptor adenylate cyclases possibly serves as the major trigger for cAMP, though the “ligands” or specific conditions for triggering such “receptors” are not defined yet. The phenotypic attributes like cell death, mitochondrial depolarization and G1 arrest illustrated in this study or social motility [26], flagellar wave reversal [14] are expected to be linked with it or some unidentified upstream effectors. Interestingly 8-pCPTcAMP is less antiproliferative to *L. donovani* compared to *T. brucei* [11,25], indicating variation of response against this cAMP-analogue between trypanosomatids. Moreover, the nonhydrolyzable analogue Sp-8-pCPTcAMPS, elicited comparable response in terms of cell cycle analysis and mitochondrial functioning. Such observations hint that possibly the anti-proliferative/ differentiation inducing action of hydrolysis product 8-pCPTcAMP as observed in *T. brucei* [27], is not conserved in *Leishmania*. However the EC50 values for the cAMP-analogues, determined in this study, showed subtle deviation across experiments, suggesting responsiveness against the analogues is delicately linked to physiological state of test *L. donovani* population. Intensive forward genetic approach combined with systemic analysis might shed light on mechanistic details of cAMP mediated responses and variations among trypanosomatids.

Though cAMP has cytoprotective effect against oxidative stress in *Leishmania*, in *Trypanosma* cAMP-phosphodiesterase inhibitor cpdA displays potent antitrypanosomal activity [25], indicating long term potentiation of cAMP triggering probably induce cell death in kinetoplastida parasites. Such observations provide genetic and pharmacological validation of using PDEs as novel drug targets for diseases caused by the kinetoplastid parasites including *Leishmania* [19]. Sebastián-Pérez et al. already projected PDE-inhibitory imidazole derivatives as candidate antileishmanial, our study corroborates PDEs as promising targets and explores the possibility of repurposing human PDE inhibitors as the starting framework for the design of evaluating in use inhibitors with substantial data for human application. This proposition is supported by the inhibitory effects of some human PDE inhibitors observed on *T.cruzi* PDEC e.g. etazolate inhibits human PDE4 and TcrPDEC with the IC_50_ values of 2 and 0.7 μm [9]. Among the PDE inhibitors used in this study, etazolate, a pyrazolopyridine derivative showed maximum anti-proliferative activity against *Leishmania* parasites with least cytotoxic effect on macrophage cells cultured *in vitro*. Further it was observed that it significantly affected the cell cycle progression and mitochondrial membrane potential of the parasite, therefore we assessed *in vitro* for their ability to clear parasite load within the macrophage cells. Though initial reports in *T. brucei* suggested that etazolate inhibits growth substantially though it does not affect intracellular cAMP pool in the parasite, a number of studies suggested that it can potentially inhibit specific PDEs from *T. cruzi* and at least two PDE from *Leishmania* [9,17]. Since etazolate apparently have noimpact of total cellular cAMP pool [5]), it is anticipated that anti-parasitic activity of etazolate is due to microdomain-specific modulation of cAMP response or by binding to some unidentified effector(s). Etazolate, primarily a PDE4 inhibitor [24,28], is a well-established drug of choice with no major side effects reported [28] in preclinical studies as well as pharmacokinetic and safety profiles in Phase I and Phase IIaclinical studies. Etazolate produced antidepressant like effects in animal models of depression and at the same time it could be used in the treatment of Alzheimer’s disease. Moreover, chronic etazolate treatment at a dose between 0.5 and 1 mg/kg treatment significantly diminished traumatic brain injury induced anomalies [29]. Earlier the drug was demonstrated to be well-tolerated and devoid of side effects by Marcade et al. (2008) based on their study on guanea pigs where they administered the drug at a dose of 10 mg/kg for 15 days [30,31]. In coherence to these reports and based on our *in vitro* cytotoxicity analysis on RAW264.7 cell line in mice, we tested dose range between 5 to 20 mg/kg which appeared to be well tolerated by the mice. With its potential inhibitory impact on amastigote proliferation *in vitro* and in experimental animal models for VL, etazolate can be considered a drug for repurposing against VL and can serve as a candidate molecule for synthesizing more selective drugs. Bearing in mind the urgent need for new therapies and the desirability of a relatively short duration of treatment, the requirement for very stringent selectivity might be relaxed somewhat if the drug was efficacious and its side effects were acceptable and reduced compared to current available treatments. Devoid of gross side effects, etazolate, as suggested by the observations can be exploited for developing intervention strategies against leishmaniasis.

## Materials and methods

### 1. Ethics Statement

The entire study was in strict accordance with the recommendations in the Guide for the Care and Use of Laboratory Animals of the Committee for the Purpose of Control and Supervision of Experiments on Animals (CPCSEA). The protocol was approved by the Institutional Animal Ethics Committee on Animal Experiments of the University of Kalyani (892/GO/Re/S/01/CPCSEA).

### 2. Parasite culture and infection

The pathogenic strain of *Leishmania donovani* AG83 (MHOM/IN/1993/Ag83) (obtained from Dr. Pijush K. Das) was maintained in BALB/c mice and promastigotes were cultured in medium 199 (M199; Invitrogen, Carlsbad, CA, USA) with Hank’s salt which contains Hepes buffer (12mM), L-glutamine (20mM), 10% heat-inactivated fetal calf serum, 50 U/ml penicillin and 50 μg/ml streptomycin. The promastigotes were isolated from infected spleens of BALB/c mice by culturing the respective spleens in M199 at 22°C for 5 days. Simultaneously, RAW 264.7 (obtained from NCCS, Pune, India), adherent murine macrophage cell line was cultured in RPMI 1640 (Invitrogen) supplemented with 10% fetal calf serum (FCS), 100 U/ml penicillin and 100 μg/ml streptomycin at 37°C with 5% CO_2_. For *in vivo* infection, BALB/c mice (Saha Enterprise, CPCSEA Regd. No: 1828/PO/Bt/S/15/CPCSEA) of approximately 20 g of body weight were injected with 1 x 10^7^ stationary phase promastigotes of *L. donovani* via the tail vein. Etazolate at a dose of 10 mg/kg/day was administered daily over a period of 30 days following 2 week of post-infection. Parasites were isolated by aseptically removing the spleen and liver of the infected mice and parasite burdens were assessed by Leishman-Donovan units (LDUs) by calculating the number of parasites per 1000 nucleated cells x organ weight (g).

### 3. Parasite viability assay

Parasite viability was measured by incubating the treated and control cells in 0.5 mg/ml 3-(4,5-dimethylthiazol-2-yl)-2, 5-diphenyltetrazolium bromide (MTT) for 3 hours followed by subsequent addition of 100 μl of 0.04 N HCl in isopropyl alcohol. The principle underlying the MTT assay is that living mitochondria convert MTT to a dark blue compound, formazan, which is soluble in acid isopropyl alcohol. Formazan was detected at 570 nm on the microplate reader. The percentage viability was obtained by calculating the ratio of OD values in wells with treated cells versus the wells with controls multiplied by 100.

### 4. Cell motility

Live *Leishmania* cells were visualized under bright field of confocal microscope on grooved slides which provides the freedom of movement to the parasites. Flagellar movements of both treated and control cells were analysed. Numerous fields were used and the distribution of counting areas was chosen in a non-biased manner along the total surface area of the cover-slip for graphical representation of the motile parasites.

### 5. Flow cytometry and cell cycle analysis

5 ml of parasite (0.5-1×10^7^ cells/ml) was centrifuged, washed twice in PBS and re-suspended in 70% ice-cold methanol for the purpose of fixation and the cells were stored at −20°C for future use. The cells were treated with 20 mg/ml of RNase A and incubated at 37°C for 1 hour before proceeding to analysis. Followed by this, propidium iodide was added for staining the DNA and 20,000 cells were analysed for DNA content using BD FACS Aria III and the distribution of G_1_, S and G_2_/M phases were then calculated from each histogram in BD FACS Diva Software.

### 6. Macrophage infectivity assay

RAW 264.7 cells were cultured in RPMI-1640 and 10% FCS, counted, centrifuged and re-suspended in culture medium and distributed over 18 mm^2^ coverslips and cultured overnight. Cells were treated with PDE inhibitors as mentioned in the result section and infected with *L. donovani* promastigotes at a ratio of 10 parasites per macrophage and the infection was allowed to proceed for overnight. For the determination of parasite number within the macrophage, cells were fixed in methanol and then stained with DAPI (4’,6-diamidino-2-phenylindole) (1 μg/ml) in PBS containing 10 μg/ml of RNase A (Sigma-Aldrich). Cells were visualized using a Olympus IX 81 microscope with FV1000 confocal system using a 100X oil immersion Plan Apo (N.A. 1.45) objective and analysed by Olympus Fluoview Software (version 3.1a, Tokyo, Japan).

### 7. BrdU incorporation assay and 1N/1K/1F enumeration assay

Promastigotes were treated with different concentrations of cAMP analogues and PDE inhibitors followed by 2 hours incubation with 100 μM of 5-bromo-2-Deoxyuridine (BrdU) as per standard protocol and the percentage BrdU incorporation was studied by BrdU cell proliferation assay (Millipore). For the determination of number and position of nucleus (N), kinetoplast (K) and flagellum (F) in both treated and non-treated cells, parasites were harvested by centrifugation at 1000 g for 10 minutes at 4°C, washed with 1x PBS and fixed on slide with methanol followed by DAPI (4’,6-diamidino-2-phenylindole) (1 μg/ml) in PBS containing 10 μg/ml of RNase A (Sigma-Aldrich) to stain nuclear and kinetoplast DNA and analysed immediately. Images were acquired using an Olympus BX61 microscope, at 1000X magnification, and an Olympus DP71 digital camera and were analysed using Image Pro Plus Software (Media Cybrernetics).

### 8. ATP bio-luminescent assay and measurement of mitochondrial membrane potential

Parasites were centrifuged, resuspended in medium and treated with cAMPanalogs and PDE inhibitors and levels of cellular ATP production was studied by ATP bio-luminescent assay (Cayman Chemicals) as per kit’s protocol.

Mitochondrial membrane potential was checked by JC-1 FACS analysis (BD Mitoscreen Mitochondrial Membrane Potential Detection Kit, BD Biosciences). Treated and non-treated cells were incubated with JC-1 for 15-20 minutes, a mitochondrial vital dye, washed with PBS and prepared for flow cytometry. Cells were studied in BD FACS Canto and BD FACS Diva software was used for studying the dot plots.

### 9. Statistical analysis

All the results shown are representatives of at least three independent experiments. Cultures were set in triplicate and the results are mean + SD. Statistical differences among the sets of data were determined by Student’s t-test with a P-value <0.05 considered to be significant.

## Acknowledgment

We would like to acknowledge DST-INSPIRE Faculty Programme, DST-PURSE Programme, University of Kalyani, DST-FIST, Department of Zoology, University of Kalyani, PRG, University of Kalyani, NASI Senior Scientist Fellowship.

## Funding

AB received grant from DST-INSPIRE Faculty Programme (IFA-12 LSBM-22), Govt. of India.

## Supporting Information Legends

**Figure S1.** Dose responsive viability assay for cAMP analogue and PDE inhibitor treatments. Log phase promastigotes were pretreated with various concentrations of 8-pCPTcAMP (A) and Sp-8-pCPTcAMPS (B); etazolate (C), dipyridamole (D), trequinsin (E), rolipram (F), EHNA (G) or zaprinast (H) for 24h followed by MTT assay. Dotted lines represent untreated promastigotes analyzed in parallel to treated population.

**Table-S1. Cell cycle progression after cAMP-analogue treatment.** G1 synchronized L. donovani promastigotes were exposed to various doses of cAMP analogues. Cell cycle stage specific distribution of the populations were analyzed by propidium iodide nuclear staining based FACS analysis after various time periods.

**Table-S2. Cell cycle progression after PDE inhibitor treatment.** G1 synchronized L. donovani promastigotes were exposed to various doses of PDE inhibitors. Cell cycle stage specific distribution of the populations were analyzed by propidium iodide nuclear staining based FACS analysis after various time periods.

